# Inner limiting Membrane Peel Extends *In vivo* Calcium Imaging of Retinal Ganglion Cell Activity Beyond the Fovea in Non-Human Primate

**DOI:** 10.1101/2024.06.02.597041

**Authors:** Hector C. Baez, Jennifer M. LaPorta, Amber D. Walker, William S. Fischer, Rachel Hollar, Sara Patterson, David A. DiLoreto, Vamsi Gullapalli, Juliette E. McGregor

## Abstract

**Purpose:** Adaptive Optics Scanning Light Ophthalmoscopy (AOSLO) paired with intravitreal injection of a viral vector coding for the calcium indicator GCaMP has enabled visualization of neuronal activity in retinal ganglion cells (RGCs) at single cell resolution in the living eye. However, the inner limiting membrane (ILM) restricts viral transduction to the fovea in humans and non-human primates (NHP), hindering both therapeutic intervention and physiological study of the retina. To address this, we explored peeling the ILM before intravitreal injection to expand calcium imaging beyond the fovea in the living primate eye.

**Methods:** Five eyes from Macaca fascicularis (age 3-10; n=3; 2 males, 1 female) underwent vitrectomy and ILM peel centered on the fovea prior to intravitreal delivery of 7m8:SNCG:GCaMP8s. RGC responses to visual flicker were evaluated using AOSLO calcium imaging 1-6 months post intravitreal injection.

**Results:** Calcium activity was observed in RGCs throughout the ILM peeled area in all eyes, representing a mean 8-fold increase in accessible recording area relative to a representative control eye. RGC responses in the ILM peeled and control eyes were comparable and showed no significant decrease over the 6 months following the procedure. In addition, we demonstrated that activity can be recorded directly from the retinal nerve fiber layer.

**Conclusions:** Peeling the ILM is a viable strategy to expand viral access to the GCL for gene therapies in NHP. Overall, this approach has potential to advance visual neuroscience, including pre-clinical evaluation of retinal function, detection of vision loss, and assessment of therapeutic interventions.

## Introduction

The retina is a highly specialized tissue in the back of the eye that plays a critical role in the capture and processing of visual information. Advances in methods to study retinal function have enhanced our ability to understand retinal physiology, disease and to evaluate and optimize therapeutic interventions. In vivo measures of function allow us to gain insight into retinal activity in an intact tissue with the integrity of both circuitry and bodily systems maintained. This is particularly important for pre-clinical development where the immune system may be provoked by the intervention. In vivo recording of retinal function also enables indefinite longitudinal follow up. The non-human primate (NHP) is the gold standard animal model for both retinal physiological study and pre-clinical testing of novel therapies, owing to the close similarity between the visual and immune systems of humans. In vivo measures of function are therefore particularly appealing in NHP where numbers are few but the potential for translation is high.

The ideal *in vivo* functional testing paradigm would evaluate retinal activity at the cellular scale throughout the retina. Multi-electrode array and patch clamp electrophysiology in retinal explants are widely used and allow recordings with high temporal resolution, but rapid degradation of the explant precludes longitudinal measurements in a single individual, complicating assessment of pathologies or therapies particularly when the time course is uncertain ^1,2^. Electroretinography (ERG) enables repeated non-invasive measurement of retinal responses in vivo ^3,4^, but does not read out activity at the cellular scale. Behavioral testing can complement optical and electrophysiological ^5^, but does not provide direct insight into function at the retinal level or at the cellular scale.

By pairing adaptive optics scanning laser ophthalmoscopy (AOSLO) which enables high resolution retinal imaging ^6^ with viral vector mediated expression of the genetically encoded calcium indicator GCaMP, it is now possible to optically read out RGC activity at the cellular scale in NHP *in vivo* ^7–9^. However, viral transduction of RGCs following intravitreal injection of an AAV has previously been spatially restricted to a ring of RGCs serving the most central foveal cones in primates ^10^ as the inner limiting membrane (ILM), a thin border between the retina and vitreous fluid formed by astrocytes and end feet of Müller cells, creates a barrier to viral penetration outside of the foveal pit ^11^. Peeling the ILM prior to intravitreal injection has reliably expanded viral transduction of GFP in retinal neurons ^12–14^ and this approach has been proposed as a strategy to dramatically expand the area of RGCs transduced by gene therapies.

In this study we dramatically expand the area of the NHP retina accessible for optical recording of function in vivo by removing the ILM prior to intravitreal injection of the viral vector coding for GCaMP. Not only does this expand the potential utility of this optical recording platform in vivo but it also provides insight into the impact of the ILM procedure on the retina at the cellular scale in the living eye. We obtain RGC responses at eccentricities that were previously inaccessible in multiple animals and present the first optical recordings from the retinal nerve fiber layer (RNFL). Longitudinal calcium imaging AOSLO is performed to together with OCT and fundus photography to better understand outcomes associated with the ILM peel procedure.

## Materials and Methods

### Experimental Design

Five experimental eyes and one control eye from 3 Macaca fascicularis (2 male, one female) were used in this study. All five experimental eyes were vitrectomized and underwent peeling of the ILM centered on the fovea (Figure 1A-B). Six to eight weeks later, intravitreal injection of a viral vector was performed to produce GCaMP expression in the ganglion cell layer (Figure 1C). Three of the five experimental eyes received intravitreal injection of viral vector AAV2.7m8-SNCG-GCaMP8s while the two eyes of macaque M3 received the GCaMP8m variant. One control eye did not undergo vitrectomy or ILM peeling (Table 1). We did not perform a vitrectomy without ILM peel as an additional control because previous studies have established that vitrectomy alone insufficient to extend viral transduction ^12^.Scanning laser ophthalmoscopy (cSLO) confirmed the presence of GCaMP in the area corresponding to region of ILM peeling surgical peel (Figure 1D). cSLO, OCT, and color fundus photography were performed longitudinally up to six months post procedure. Functional assessment at the cellular scale *in vivo* was conducted using fluorescence AOSLO in combination with a red LED flashing stimulus (Figure 1E-G). Retinal imaging was conducted between the fovea and the ILM peel edge in all meridians.

**Figure 1:**
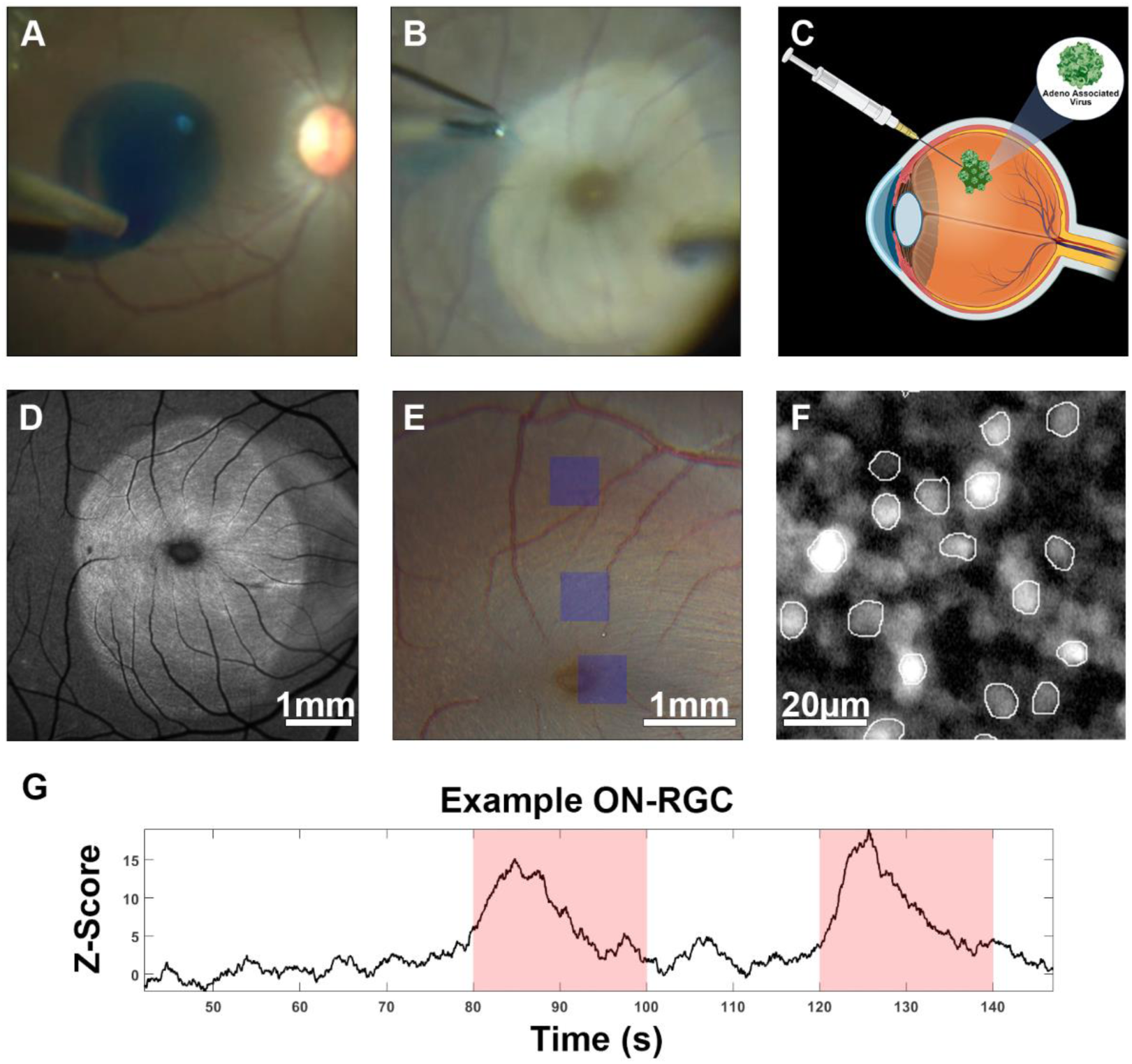
Overview of study. **A.** Following a pars plana vitrectomy, the ILM is stained with brilliant blue dye. **B.** Mechanical force lifts the ILM in a circular pattern around the fovea using forceps. **C.** Intravitreal injection delivers viral vector to the retina. **D.** A blue autofluorescence (BAF) confocal scanning laser ophthalmoscope (cSLO) confirmed the presence of GCaMP in a pattern matching the surgical peel two months post injection. **E.** Functional assessments were conducted at retinal imaging locations using a blue 488nm excitation source in combination with a red LED flashing stimulus. **F.** Individual RGCs were segmented and **G.** their responses to the LED stimulus tabulated.

**Table 1:** Summary of demographic and experimental parameters for all eyes involved in the study. The sex and age of the primate, immune suppression status quantity and type of viral vector injected intravitreally, and the time elapsing between the ILM peel and the intravitreal injection in days is listed for each eye.

| *Macaque (eye)* | *sex* | *Age at time of ILM peel* | *Inactivating antibodies to AAV2* | *Time between immune suppression and ILM peel (days)* | *Time between ILM peel and viral vector Injection (days)* | *Time of AAV2 injection after initial injection in fellow eye (days)* | *GCaMP Variant* |
| --- | --- | --- | --- | --- | --- | --- | --- |
| *M1 (OS)* | *M* | Control | 1:4 | Control | Control | --- | GCaMP8s |
| *M1 (OD)* |  | 10y 9m |  | 31 | 28 | 29 | GCaMP8s |
| *M2 (OS)* | *M* | 4y 2m | 1:5 | 27 | 50 | 41 | GCaMP8s |
| *M2 (OD)* |  | 4y 3m |  | 62 | 55 | --- | GCaMP8s |
| *M3 (OS)* | *F* | 3y 10m | 1:10 | 98 | 42 | 35 | GCaMP8m |
| *M3 (OD)* |  | 3y 11m |  | 126 | 48 | --- | GCaMP8m |

### Animal welfare

The macaques used in this study were housed in pairs in an AAALAC accredited facility in compliance with the University of Rochester University Committee on Animal Resources (UCAR). They had free access to water and primate biscuits, providing a complete and nutritious diet. To supplement this, they were given various treats multiple times daily, such as nuts, dried fruit, vitamin chews, and a large variety of fresh fruits and vegetables. They were also given “high value” treats during training, including Ensure, marshmallow snacks, and fruit gummies. An animal behaviorist and Veterinary Technicians provided novel enrichment items multiple times a week, including items such as grass, oranges with peels still on, treat filled forage boxes, grapevines, water play, and popcorn balls. Daily enrichment included mirrors, puzzle feeders, durable chew toys, daily television time and/or music and regular visits from staff to offer treats and socialization. All macaques were cared for by the Department of Comparative Medicine veterinary staff, including three full-time Veterinarians, five Veterinary Technicians, and an animal care staff who monitored animal health and comfort at least three times daily. The Animal Behaviorist regularly recorded and analyzed primate behavior to monitor any signs of stress. This study was carried out in strict accordance with the Association for Research in Vision and Ophthalmoscopy (ARVO) Statement for the Use of Animals and the recommendations in the Guide for the Care and Use of Laboratory Animals of the National Institutes of Health. The protocol was approved by the University Committee on Animal Resources of the University of Rochester (PHS assurance number: D16-00188(A3292-01)).

### Immunosuppression

All macaques in this experiment received subcutaneous Cyclosporine injections daily. Blood trough levels were collected to titrate the dose of 150-250 ng/mL and then maintained at that level. The timing of immune suppression relative to ILM peel surgery and viral vector injections are detailed in Table 1.

### Animal preparation for ILM peel surgery

The surgical subject was fasted overnight with free access to water. The animal was sedated using ketamine (100mg/mL) and medetomidine (20mg/mL), then intubated and kept at proper anesthetic depth via 1.5-3% isoflurane throughout the procedure. The animal was positioned in dorsal recumbency on a padded surgical table. A Bair Hugger warming system was placed over the animal to maintain body temperature. Vital signs including temperature, heart rate and rhythm, respirations, end tidal CO_2_, blood pressure, SpO_2_, and reflexes were monitored consistently and recorded every fifteen minutes. Pupil dilation was accomplished using 1% atropine ophthalmic solution. An aseptic preparation was performed on the surgical eye and surrounding area using 5% betadine solution and flushed with sterile 0.9% saline. The subject’s eye and body were then draped in a sterile fashion.

### ILM Peel procedure

The ILM peel procedures performed in NHP closely followed the protocol used in human surgery. A standard 3-port pars plana vitrectomy was performed through 25-gauge cannulas using the Constellation vitrectomy system (Alcon, Ft. Worth, TX), OPMI Lumera 700 operating microscope, and RESIGHT viewing system (Carl Zeiss Meditec AG, Jena, Germany). Triamcinolone was injected intravitreally to visualize the vitreous. A posterior vitreous detachment was induced using aspiration from the vitrector or a soft tip cannula. The peripheral vitreous was trimmed. Brilliant Blue G Ophthalmic Solution (DORC) dye was placed over the macula and the ILM stained for 2 minutes (Supplementary Movie 1). A fluid-fluid exchange was performed to remove the dye. The stained ILM was then peeled 360 degrees, centered on the fovea, in approximately 5 to 6-disc diameter area in the macula using ILM peeling forceps (Alcon, Ft. Worth, TX) (Supplementary Movie 2). The three ports were then removed at the end of the surgery and the sclerotomies sutured. The animal was given an intravitreal injection of vancomycin (1mg/0.1mL) and ceftazidime (2mg/0.1mL). Immunosuppressant Cyclosporine (150-250ng/mL), anti-inflammatory meloxicam (5mg/mL), analgesic buprenorphine SR (3mg/mL), and an anesthetic reversal of antisedan (5mg/mL) were all administered post-operatively.

### Intravitreal Injection of viral vector

Intravitreal injection of AAV2-7m8.SNCG.jGCaMP8s.WPRE.SV40 (100 µL at titer 2.62×10^13^ GC/mL), or AAV2-7m8.SNCG.jGCaMP8m.WPRE.SV40 (100 µL at titer ≥ 2.46×10^13^ GC/mL), was performed as previously described in Yin et al 2011 ^15^. Briefly, the eye was sterilized with 5% diluted betadine before the vector was injected into the vitreous through the pars plana using a tuberculin syringe and 30-gauge needle. All viral vectors were synthesized by the University of Pennsylvania Vector Core. Neutralizing antibodies to AAV2 were 1:25 or lower in all four injected animals prior to the study. Following the intravitreal injection of the vector, each eye was imaged weekly with a conventional scanning light ophthalmoscope (Heidelberg Spectralis, Heidelberg Engineering, Franklin, MA) using the 488 nm autofluorescence modality, to determine the onset of expression, image quality and to monitor eye health. The animal received 2mg/0.05mL of triamcinolone intravitreally (Kenalog-40) following the injection to treat for symptoms of uveitis.

### Animal preparation for imaging

All monkeys were fasted from 4 to 18 h prior to anesthesia induction. Anesthesia induction began with 5 to 20 mg/kg Ketamine, 0.25 mg/kg Midazolam, and 0.017 mg/kg Glycopyrrolate intramuscularly. The pupil was dilated with a combination of tropicamide 1% and phenylephrine 2.5%. In cases of minimal pupil dilation within the standard time, phenylephrine 10% and/or cyclopentolate 1% drops were administered. Flurbiprofen Ophthalmic drops are given to the target eye, to prevent any discomfort from prolonged use of an eye lid speculum. Both eyes were covered with a hydrating ophthalmic gel (GenTeal). The target eye then had the lid speculum placed to keep the eye open during imaging and a contact lens was placed to ensure corneal protection. The fellow eye was taped closed with porous tape, to protect the cornea from drying.

The animal was placed in a stereotaxic cart. Prior to intubation, an oxygen mask with 1–2% isoflurane, was placed over the monkey’s face to allow for adequate sedation for intubation. An intravenous drip of Ringer’s lactate with 5% dextrose was maintained at 5 ml/kg/hr for the duration of imaging. The monkey was intubated and maintained at a surgical plane of anesthesia with isoflurane 1.0–2.5%. A Bair Hugger warming system was placed over the monkey to maintain body temperature. Monitoring was conducted as previously described.

After a surgical plane of anesthesia had been established, a Rocuronium infusion (800mcg/ml) was given with a dose range from 200-600mcg/kg/hr, after an initial bolus dose of 100-400mcg/kg. Rocuronium is administered to effect, determined by eye movement, and the lowest effective dose per animal is used. Once respirations ceased, the monkey was maintained on a ventilator until imaging was over and the infusion was turned off. Once a peripheral nerve response was established, an intravenous dose of glycopyrrolate 0.01 mg/kg was given. Five minutes after the glycopyrrolate, neostigmine 0.05 mg/kg was given intravenously. The monkey was monitored for indications of breathing against the ventilator and then removed from the ventilator once able to breath without assistance. The monkey was removed from the isoflurane no sooner than fifteen minutes after the neostigmine injection to ensure stability off the ventilator. The monkey was then allowed to wake up and extubated once all reflexes had returned.

### Clinical imaging

Confocal scanning light ophthalmoscopy (cSLO), optical coherence tomography (OCT) and color fundus photography were performed to pre- and post-procedure using a commercially available Spectralis HRA (Heidelberg Engineering) and fundus camera (Topcon 50EX). The blue autofluorescence modality on the spectralis HRA was used to capture GCaMP fluorescence using the 488nm excitation wavelength and a longpass filter at 500nm. GCaMP fluorescence was also captured using the fundus camera equipped with filters selective for GCaMP excitation and emission (ex. 472/30 nm em. 520/28 nm). 30- and 55-degree OCT images centered on the fovea were collected pre- and post-ILM peel, pre-and post-intravitreal injection, and roughly every 30 days thereafter. OCT data from the optic nerve head in addition to slit lamp photographs (Zeiss) were recorded at the same timepoints to document the time course and severity of any inflammation.

### Assessing ocular inflammation

A modified Hackett-McDonald scoring system ^16^ was used to evaluate ocular inflammation. A Veterinary technician evaluated the pupil reflex and conjunctival discharge prior to pupil dilation. A comprehensive set of slit lamp images were recorded, randomized and graded by an ophthalmologist (VG). The scoring system is available in the supplementary materials.

### Calcium Imaging Adaptive Optics Scanning Laser Ophthalmoscopy

AOSLO data were collected using a previously described AOSLO instrument ^17,18^. An 847nm diode laser (QPhotonics, 100-136uW/mm2) was used as a wavefront sensing beacon in combination with a Shack-Hartman wavefront sensor and a deformable mirror (ALPAO) for measuring and correcting the optical aberrations of the eye. A 796nm superluminescent diode (Superlum, 700-850uW/mm^2^) was used to image the cone photoreceptor layer using a 2 airy disc (20um) pinhole to collect the reflected light. Simultaneously, a 488nm laser (Qioptiq, 57uW/mm2) focused on the RGC layer was used to excite GCaMP and emitted fluorescence was collected using a 7.5 airy disc (75um) pinhole. Laser powers were calibrated prior to every imaging session to ensure consistency across sessions. To correct for longitudinal chromatic aberration between the 796nm source and 488nm sources on the detector, motorized stages were adjusted to optimize their positions *in vivo* prior to each experiment. The adaptive optics control facilitated fine focus on the retinal ganglion layer of the 2.54° by 2.54° field of view at each retinal location. GCaMP fluorescence (emission 535/20 nm) was recorded in response to 7 degree uniform visual stimulus presented through a Maxwellian view system ^19^. A Thorlabs mounted LED with a 661/20 nm emission filter was used as the visual stimulus to avoid the GCaMP excitation spectrum and consisted of 20 second contrast increments and decrements against a mean light level (3 μW/mm^2^). Trials began with a 60 second system background period with no laser emission, this was followed by a 120 second light adaptation period to the 488nm imaging laser and the mean luminance of the visual stimulus before 20 second increments and decrements began. Fluorescence was recorded during the system background, period of constant illumination and the stimulation period. Each trial was repeated twice at each retinal location. Total exposure to all laser sources was kept below the Rochester exposure limit ^20^, designed to be two-fold more conservative than the ANSI standard ^21^.

### Data Analysis

Adaptive optics calcium imaging was stored as an uncompressed AVI video format. A frame-by-frame stimulus information was written to a csv file, providing an accurate record for analysis. The reflectance videos of the photoreceptor mosaic were registered offline using a strip-based cross correlation algorithm ^22^, that corrected translational motion between frames and intraframe warping due retinal motion. Each frame of the GCaMP fluorescence video was co-registered to the photoreceptor reflectance video. The frames of the fluorescence video were temporally integrated to form a high SNR z projection image that was used to create an RGC segmentation mask using FIJI. RGC segmentation masks were then imported into MATLAB and used to tabulate individual cell traces for each imaging location. These cell traces were then filtered using a 50-frame moving median filter. The Z-score was calculated as the GCaMP response amplitude (signal) minus the mean amplitude of the baseline adapted response (noise), all divided by the standard deviation of the noise. The criterion for a responsive cell was set to 2.5 standard deviations above the noise. Statistical analyses were performed using MATLAB two-sample T-test, with group spread reported as mean ± SD. Fluorescence cSLO images were scaled and manually segmented using FIJI to determine the boundaries of GCaMP expression.

Following RGC imaging, we recorded from the RNFL near the optic disc in the same sessions. We used the same imaging and stimulation paradigm to collect this data with the expectation that the 7-degree uniform visual stimulus would excite ganglion cells within the area of GCaMP expression. We integrate a large area encompassing multiple passing fibers to enable extraction of population-level responses.

### Code and Data Accessibility

All code used for analysis and corresponding filtered RGC recordings will be made available on our GitHub and the open science framework repository upon publication. Raw video data is available upon request.

## Results

### ILM peel enables functional recording from RGCs at previously inaccessible eccentricities *in vivo*

To assess the impact of ILM peel on the spatial extent of GCaMP transduction in the GCL we directly compared the pattern of expression in the ILM peeled eye with the fellow eye of the same animal (M1), which received an intravitreal injection but no ILM peel (control). Figs 2A and 2B show a dramatic 5-fold increase in the area of GCaMP expressing RGCs in the ILM peeled eye (12.9 mm^2^) relative to the control (2.5 mm^2^). In the ILM peeled eye, RGCs were transduced with GCaMP out to 9.2 degrees superior to the fovea whereas in the control eye the maximum eccentricity was just 3.5 degrees. The increased fluorescence relative to the autofluorescent background observed in Figure 2B extended to the limit of the ILM peel identified by retinal landmarks in the surgical video. Furthermore, imaging with a fluorescence fundus camera equipped with filters selective for GCaMP excitation and emission confirmed that the elevated fluorescence was the extrinsic fluorophore GCaMP.

**Figure 2:**
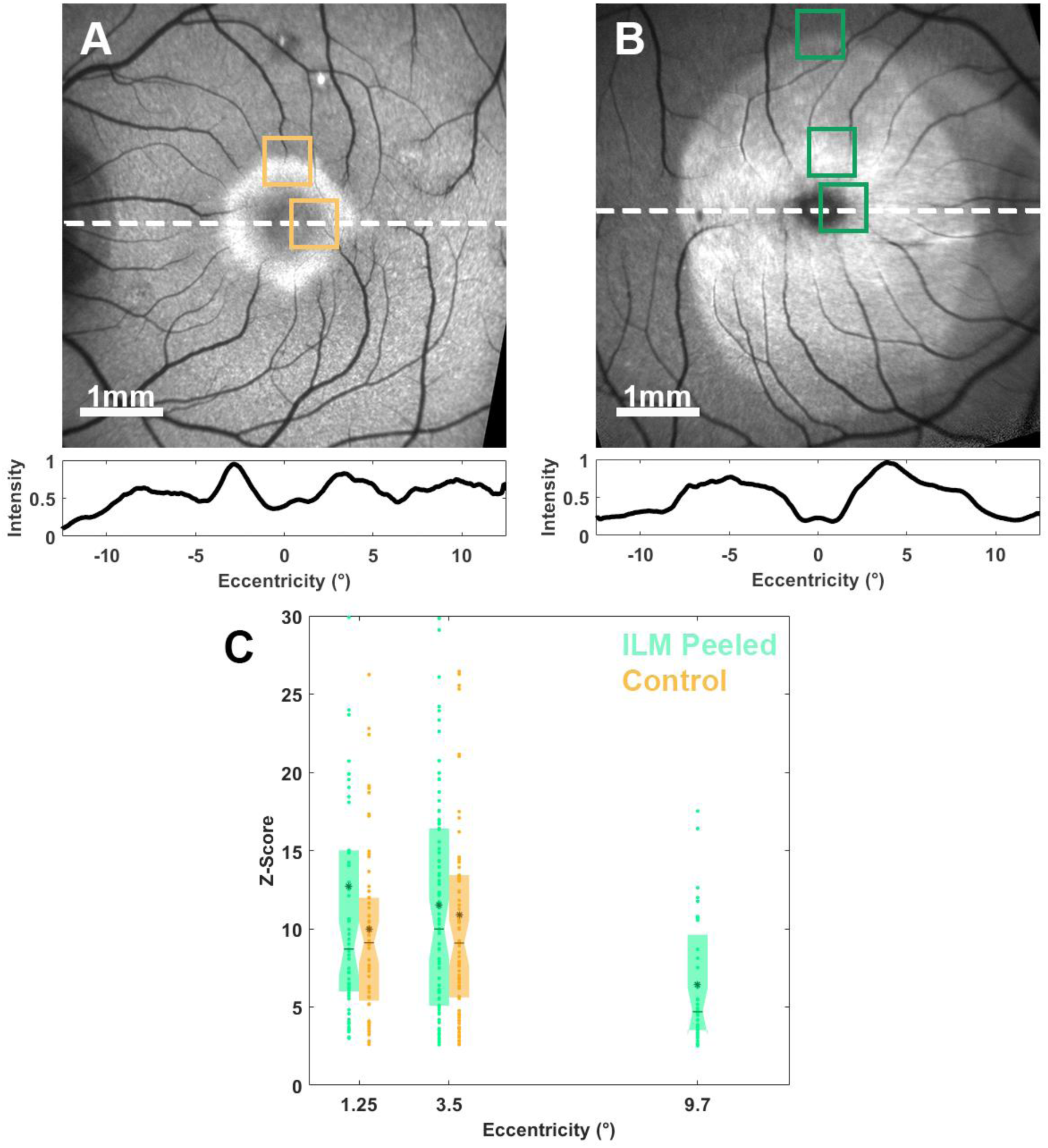
ILM Peeling enables AAV-mediated fluorescence beyond the fovea in macaque. **A.** 30°cSLO BAF images centered on the fovea, taken in each eye of M1 including the A control eye OS and **B.** the ILM peeled eye OD. GCAMP Fluorescence intensity contour (visualized in lower panel, line denoted by white dashed line) is normalized to max value and smoothed. AOSLO Imaging locations are labeled for each eye. **C.** Box plot of individual RGC Z-scores at each imaging location, grouped by retinal eccentricity. Group responses are 12.7 ± 10.4, n=67; 10.0 ± 6.6, n=62; 11.52 ± 7.5, n = 85; 10.9 ± 7.6; n = 73; 6.4 ± 3.9, n=40, respectively. Taken at 17 weeks post intravitreal injection in OS and 21 weeks OD.

To assess the responses of RGCs in the ILM peeled and control eye we performed calcium imaging AOSLO while the retina received periodic visual stimulation at 0.025Hz. Figure 1G shows an example fluorescence trace from a single ON cell in the ILM peeled eye. The magnitudes of single cell responses recorded in the ILM peeled and control eyes are shown in Figure 2C, represented as z-scores. The distribution of z-scores from individual RGCs in the ILM peeled and control eyes were not significantly different at 1° (P=0.07) or 3.5° (P=0.61), suggesting that the ILM peel did not adversely impact RGC physiology. Importantly only in the ILM peeled eye was it possible to obtain functional recording from RGCs at eccentricities beyond 3.5 degrees. Figure 2C shows responses collected from RGCs at 9.2 degrees from the foveal center. This is the first-time *in vivo* calcium imaging has been performed outside the parafovea. No RGCs fluoresced beyond four degrees from the fovea in M1 OS. Z-scores from M1 OD at 9.2 degrees were found to be significantly lower than those at 3.5 degrees in the same eye (P=0.0004).

As functional recordings were obtained *in vivo,* it was possible to return to the same region of the GCL to longitudinally monitor RGC responses over a period of months (Figure 3). For M1 OD, RGC response amplitudes at 9 degrees did not significantly change over time (Figure 3A-B; paired t test; P=0.86, P=0.42), but the amplitudes at 3 degrees significantly increased after 5 months (Figure 3C-D; paired t test; P=0.02). Compared to the non-ILM peeled fellow eye at 3.5 degrees (Figure 3E-F) that was injected first, OD had significantly higher amplitudes (paired t test; p=0.021). While responses varied week to week, no long-term decline in the calcium responses of RGCs were observed, suggesting that RGC function was not significantly compromised by the ILM peel procedure.

**Figure 3:**
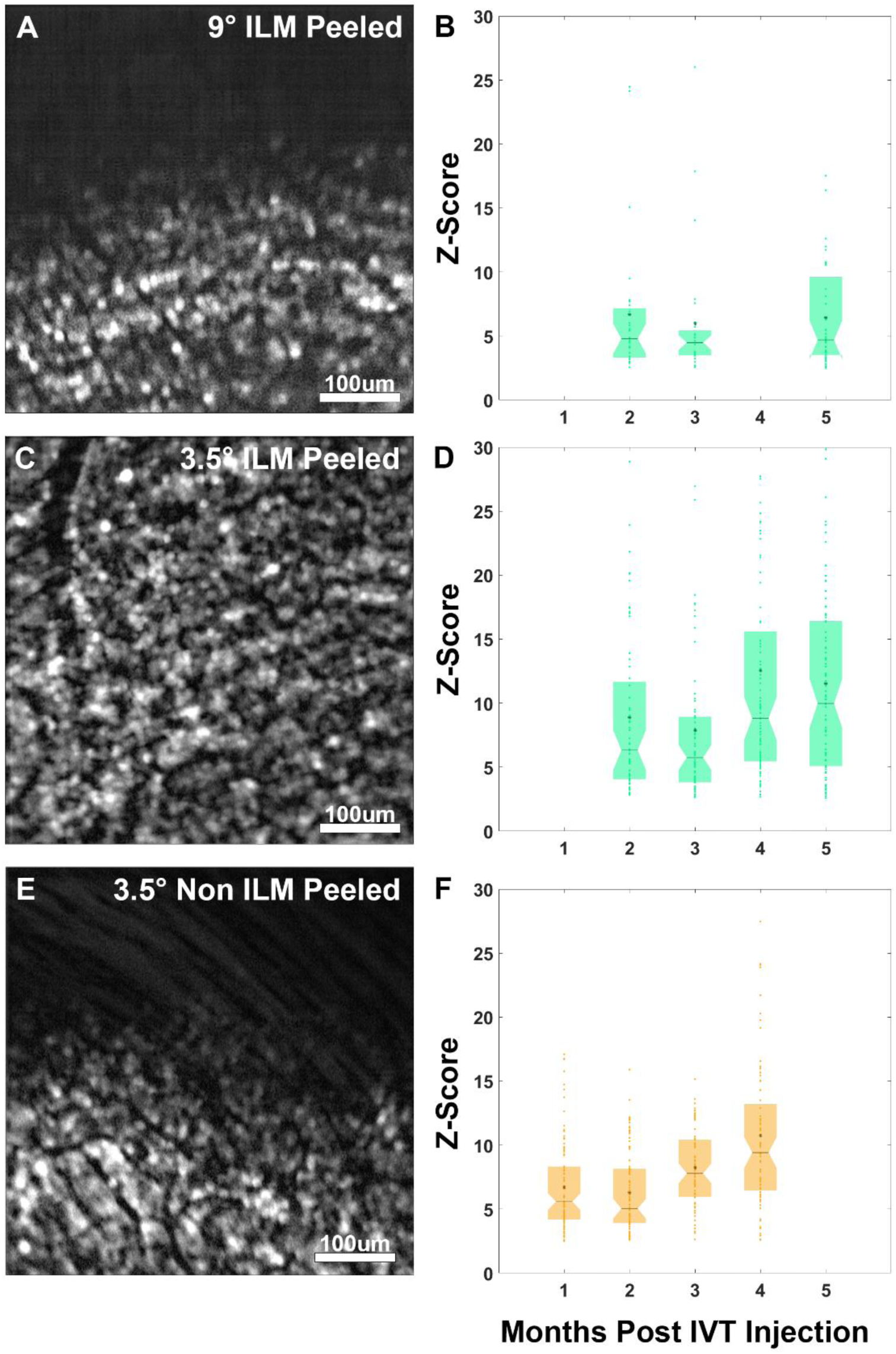
RGC remain responsive for at least 6 months after ILM peel. **A**. AOSLO GCaMP time-integrated image used to create segmentation masks at the edge of the ILM peeled area at 9° from the fovea of M1 OD. **B.** Responses at two (6.7 ± 5.7, n=27), three (6.0 ± 5.2, n=28), and five (6.4 ± 4.0, n=40) months post intravitreal injection, or three, four-, and six-months post ILM peel. **C.** AOSLO GCaMP image within the ILM peeled area at 3.5° from the fovea of M1 OD. **D.** Responses at three (8.9 ± 6.2, n=59), four (7.8 ± 6.6, n=68), five (12.5 ± 10.2, n=87), and six (11.5 ± 7.5, n=76) months post intravitreal injection. **E.** AOSLO GCaMP image of the non-peeled eye at 3.5° from the fovea of M1 OS. **F.** Responses at one (6.7 ± 3.5, n=97), two (6.3 ± 3.2, n=74), three (8.2 ± 2.9, n=81), and four (10.7 ± 6.0, n=75) months post intravitreal injection.

To assess the reproducibility of this approach an additional 4 eyes underwent ILM peeling prior to intravitreal injection of the viral vector and follow up imaging. Across all eyes we observed an average 8-fold increase in area (12.9mm^2^, 24.4 mm^2^, 20.1 mm^2^,22.7 mm^2^, 20.7 mm^2^, respectively) of GCaMP expression across all eyes relative to the control eye (2.5mm^2^) as measured using BAF cSLO (Figure 4). We extracted functional activity from RGCs in all eyes at eccentricities out to 9.2, 7.4, 7.9, 11, and 6.2 degrees respectively. There was a significant decrease in response magnitude with eccentricity in M1 OD (p=0.0003) but not in M2 OS (p=0.598), M2 OD (p=0.147), M3 OS (p=0.055), and M3 OD (p=0.122). We note that M2 had naturally occurring vessel tortuosity.

**Figure 4:**
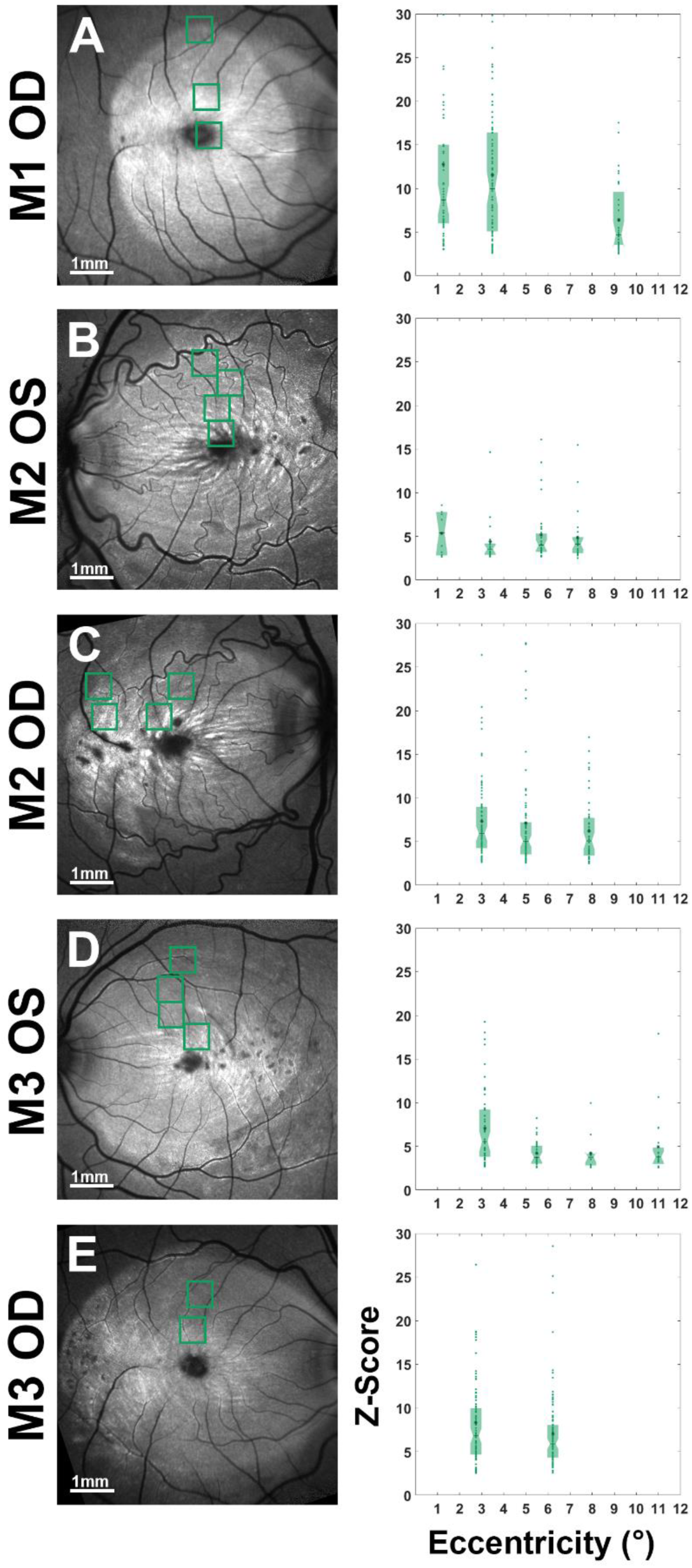
RGC response amplitudes measured in all eyes. **A.** (left) 30°cSLO BAF images centered on the fovea for all experimental eyes, imaging locations shown with blue squares. (right) Individual RGC Z-scores in box plots for M2 OD (12.7 ± 10.4,n=67, 11.5 ± 7.5,n=85, and 6.4 ± 3.95,n=40) taken five months post IVT injection, **B.** M2 OS (5.4 ± 2.5,n=12, 4.4 ± 2.9,n=17, 5.2 ± 3.3,n=30, and 4.4 ± 3.0,n=24) taken three months post IVT injection, **C.** M2 OD (7.3 ± 4.6,n=81, 7.1 ± 6.2,n=77, and 6.2 ± 3.7,n=48) taken five months post IVT injection, **D.** M3 OS (7.1 ± 4.3,n=52, 4.2 ± 1.5,n=31, 4.2 ± 2.1,n=12, and 4.9 ± 3.6,n=20) taken five months post IVT injection, **E.** M3 OD (8.3 ± 6.4,n=92, and 7.1 ± 4.4,n=99) taken six months post IVT injection.

### Optical readout of RGC activity is possible from the retinal nerve fiber layer

For the first time we show that it is possible to record not only from RGC somas but also from the RNFL near the optic disc (Figure 5). We evaluated calcium responses from the RNFL in both the ILM peeled and control eye. The investigated locations were outside of the ILM peeled area and as a result lacked fluorescing RGC somas but contained GCaMP expressing axonal fibers (Figures A,C). The averaged signal from the integrated region (Figures 5B,D) corresponded to an ‘OFF’ type response in both cases, but in the control eye the magnitude of the response was reduced by half (3.3 ± 0.02, and 1.4 ± 0.02 Z-score, respectively) (Figure 6E). In a representative location with both soma and RNFL, the RGC somas were more response than segmented RNFL bundles (10.9 ± 7.63 and 5.28 ± 3.3 Z-score, respectively) with 400-fold greater average pixel brightness.

**Figure 5:**
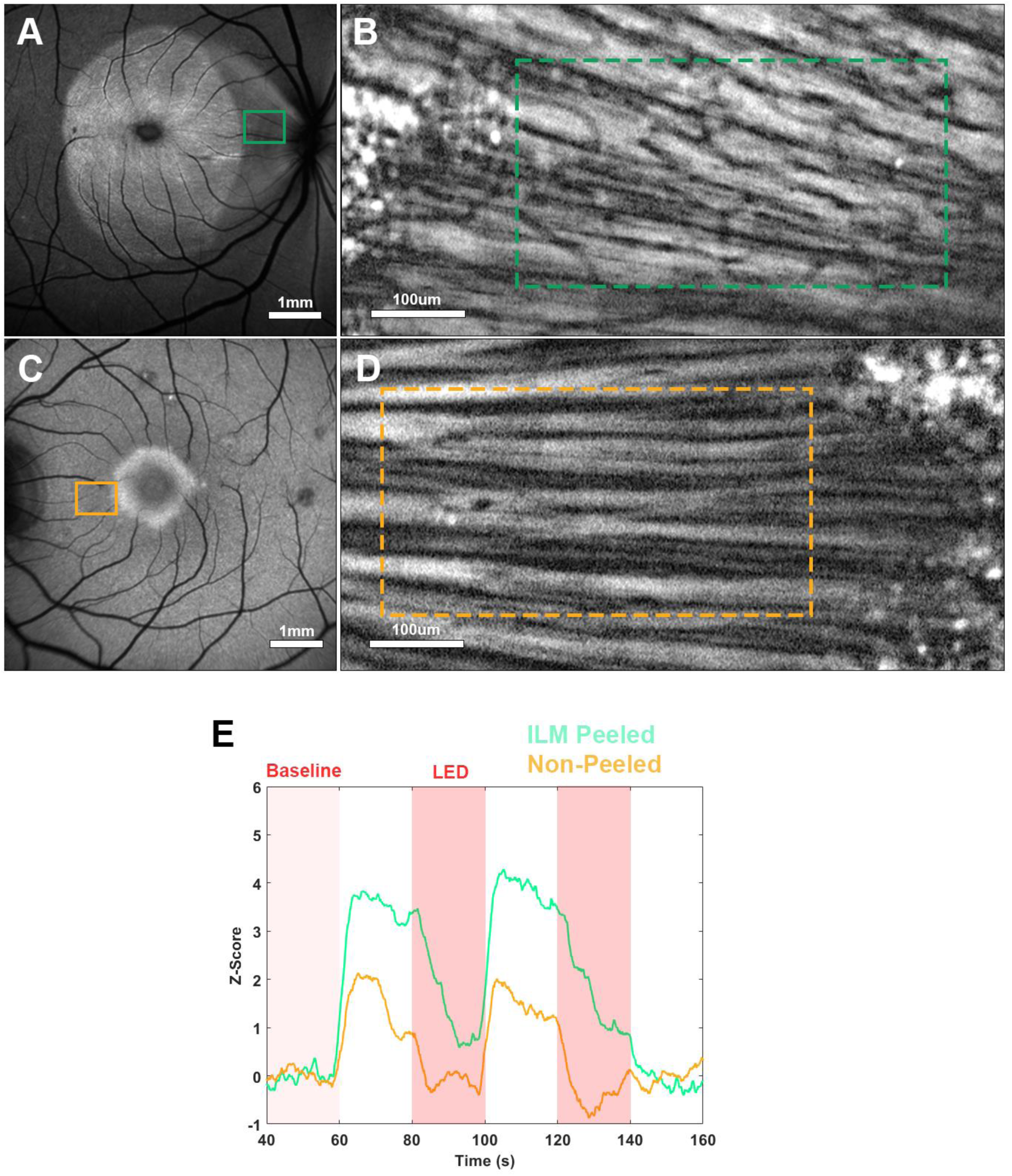
Optical readout of RGC activity from the retinal nerve fiber layer. **A.** 30°cSLO BAF images centered on the fovea of M1 OD (ILM-peeled). **B.** AOSLO GCaMP time-integrated image taken at the Nasal region shown in M1 OD. The green dotted line represents an area-integration used to construct a temporal signal for these RNFL bundles **C.** 30°cSLO BAF images centered on the fovea of M1 OS (non-peeled). **D.** The orange dotted line represents an area-integration used to construct a temporal signal for these RNFL bundles **E.** RNFL area-integrated traces responding to an OFF-onset of the LED stimulus at 60s and 100s (Z-Score responses 3.36 ± 0.02, and 1.45 ± 0.02, respectively), filtered using moving median filter as previously described.

**Figure 6:**
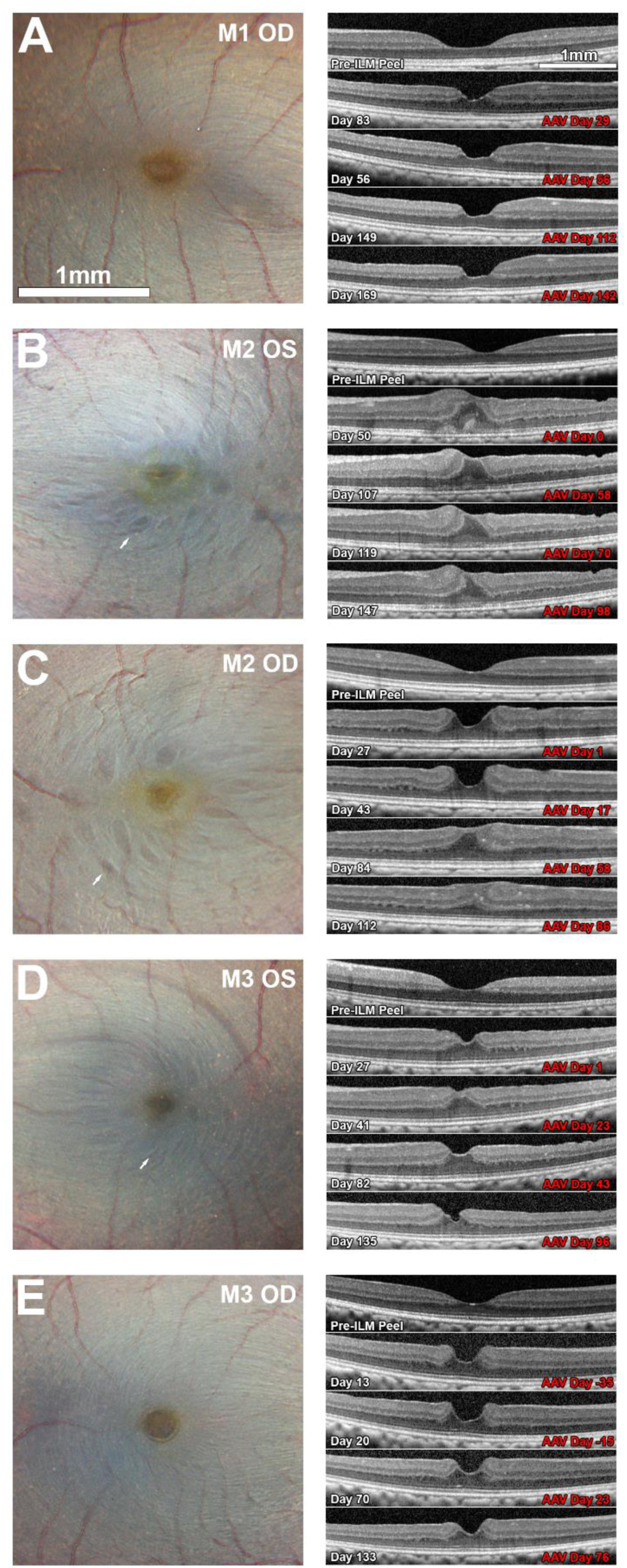
Structural imaging reveals minor morphological changes following ILM peel. **A** (left) 55° Color fundus images of all ILM peeled eyes centered on the fovea and taken two-months post ILM peel. White arrows indicate areas consistent with concentric macular dark spots. (Right) B-scan OCT of the foveal pit between 2-24 weeks post ILM peel procedure and 1-142 days post intravitreal injection.

### Structural imaging reveals minor morphological changes following ILM peel

To ensure we had a full understanding of any structural changes in the retina resulting from the ILM peel, we closely monitored all eyes over time with fundus photography and OCT (Figure 6). Alteration of the foveal contour was visible in the B-scan OCT images of all subjects, with the extent of this alteration varying between individuals. The foveal pit diameter of M1 OD (Figure 6A) and M3 OD (Figure 6E) appears slightly reduced compared to pre-ILM peel, while, the foveal contours of M2 (Figure 6B-C) and M3 OS (Figure 6D) are more dramatically altered. These changes in foveal contour persist throughout the observation period (6 months post ILM peel). No macular holes were created, but disruption to the photoreceptor layer was noted in M2 OS and improved over time.

Concentric macular dark spots (CMDS) ^23^, surrounding the fovea and aligned with the sweep of retinal nerve fibers toward the optic disc were observed in the color fundus images of 3 of the 5 eyes (Figures 6B,C,D, white arrows). Shallow dimples in the RNFL were observed post ILM peel in OCT-B scan data from all animals, consistent with previous reports of disassociated optic nerve fiber layer (DONFL) ^24,25^.

To further investigate potential damage to the GCL or RNFL, we used high resolution imaging to investigate retinal locations containing both darkened spots in color fundus (Figure 7B,D,F) and DONFL in the OCT-B scan. Our findings confirmed for the first time that dimples present in the RNFL correlated to the absence of small (< 100um) pockets of RGCs (Figure 7A,C,E). This RGC loss was variable between animals. Animal M3 OS (Figure 7D, left panel) had only one hypofluorescent region near the edge of expression (Figure 7D, right panel) while other eyes had them close to the fovea predominantly in temporal retina. To test the connection between CMDS and the RNFL dimples in OCT we investigated patches of CMDS with high resolution imaging, finding that these were not associated with missing RGCs. As one example, in M3 OS (Figure 7F-G) the DONFL region does not appear on the OCT as a dimple in the RNFL nor in the AOSLO modality as hypofluorescent.

**Figure 7:**
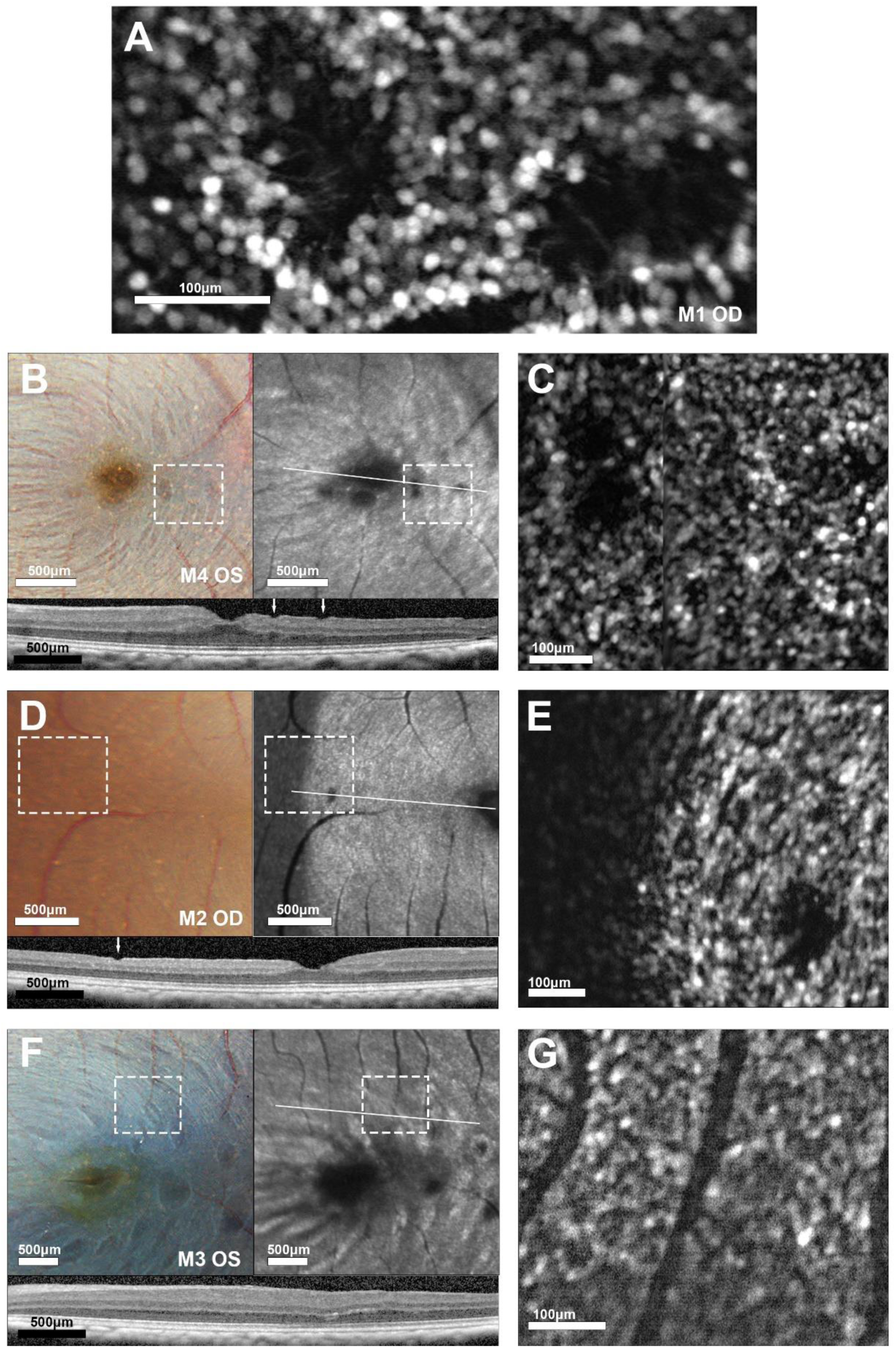
Circular retinal defects correspond to absence of ganglion cell fluorescence. **A.** AOSLO calcium image of two exemplary hypofluorescent areas in M3 OS. Dendrites are visible in this region. **B.** Color fundus (top-left) and cSLO BAF (top-right) of M3 OS showing two hypofluorescent areas that are investigated with OCT (bottom). **C.** Montaged fluorescence AOSLO image reveals dimples in OCT correspond to the two areas in AOSLO. **D.** Color fundus location with no macular dark spots but a hypofluorescent region in the cSLO BAF **E.** OCT, and AOSLO of the hyperfluorescent region seen in D. **F.** Color fundus location with CMDS streaks but no visible RNFL dimples in OCT or **G.** hypofluorescent RGC expression in AOSLO.

A board-certified ophthalmologist evaluated ocular inflammation for 4 eyes of 2 animals following intravitreal injection of the viral vector based on a modified Hackett-McDonald scoring system (Table 2) [29]. In M2, inflammation was characterized principally by conjunctival congestion, swelling, and decreased corneal transparency with pigment noted on the anterior capsule of OS up to 12 weeks post injection. M3 primarily experienced decreased corneal transparency along with minor conjunctival congestion and swelling up to 7 weeks post injection. Despite temporary inflammation, GcAMP was present in all eyes for the duration of this study.

**Table 2:** Ocular Inflammation scores. Inflammation following intravitreal injection of the viral vector in all animals was scored longitudinally by a masked board certified ophthalmologist based on a modified Hackett-McDonald scoring system ^16^. The assessment included slit lamp examination and observations of pupillary response and dilation. Scores shown are integrated from the 9 numerical measures listed in the attached Modified Hackett scoring sheet.

|  | Weeks post injection |  |  |  |  |  |  |  |  |  |  |  |  |  |  |  |  |
| --- | --- | --- | --- | --- | --- | --- | --- | --- | --- | --- | --- | --- | --- | --- | --- | --- | --- |
| Animal | -4 | -3 | -2 | -1 | 0 | 1 | 2 | 3 | 4 | 5 | 6 | 7 | 8 | 9 | 10 | 11 | 12 |
| M2 OS |  |  |  |  |  |  | 11 |  | 4 |  | 3 | 3 | 4 |  | 2 |  | 2 |
| M2 OD | 2 |  | 4 |  |  | 8 | 2 |  | 3 |  | 2 |  |  |  |  | 0 |  |
| M3 OS |  | 2 |  |  |  | 0 | 4 | 4 | 2 | 3 |  | 2 |  |  |  |  |  |
| M3 OD | 2 |  |  |  |  | 3 | 2 | 2 |  | 2 |  |  |  |  |  |  |  |

## Discussion

Surgical removal of the ILM prior to intravitreal injection of a viral vector coding for GCaMP enabled a mean 8-fold expansion of the area of the retina accessible for optical recording from RGCs across the 5 primates tested in this study. These are the first optical measurements at eccentricities beyond a small (< 3 degree) foveal ring that have been achieved *in vivo* and the first demonstration that optical recording from the RNFL is possible. Longitudinal recordings of RGC calcium responses over weeks and months showed variation but no significant long-term decline, suggesting that RGC function was not significantly compromised by the ILM peel procedure. This result is promising for future applications of ILM peeling in gene and cell therapies. In addition, we used AOSLO to investigate previously reported structural defects associated with the ILM peel procedure and observed a variable number of small hypo fluorescent regions of RGC loss, more common in temporal retina and visible as depressions in the OCT images of the GCL. This minimal loss of discrete patches of RGCs may be an acceptable tradeoff given the expanded region accessible for calcium imaging *in vivo* or for therapeutic applications.

The advances presented in this paper have implications for both retinal neuroscience, pre-clinical assessment of retinal function, and for the delivery of new therapies that could utilize the ILM peel procedure. The ability to functionally survey large areas of the GCL beyond the fovea both in vivo and at the cellular scale could be valuable in identifying and interrogating rare classes of primate RGCs and understanding how these cells change with eccentricity. From a pre-clinical standpoint, the cellular scale assessment of retinal function in vivo has potential to advance our ability to characterize disease models and detect and track restored RGC function that might be spatially and temporally sporadic, for example in the development of cell therapies. In this use case, peeling the ILM would expand the available surface area for therapeutic investigation 8-fold. Lastly surgical peel of the ILM may be a viable strategy to expand region of the retina that can be transduced by gene therapies or receive RGC transplants. The results presented here suggest that RGC function is not severely impacted by this intervention in the months after surgery and in the case of e.g. optogenetic gene therapy targeting RGCs, ILM peel could substantially increase the patients restored visual angle and any potential benefit derived from this therapy.

### Variability in RGC responses between subjects

We observed a substantial variability in Z scores between animals, with subject M1 responding with significantly higher z-scores relative to other experimental eyes. A variety of factors could be behind this, but we note that M1 was significantly older (10 as opposed to 4 years) and had fewer inactivating antibodies to AAV2 (Table 1). Age can have an impact on innate immunity, and could have affected the efficacy of viral transduction or the degree of inflammation impacting anterior segment optics. M1 also had a much shorter time between ILM peel and the first intravitreal injection, minimizing any potential obstruction of viral access to the GCL by glia mediated scar formation, as while the ILM does not regrow, Muller glia produce a basement membrane that covers most of the exposed retina ^26^.

### Functional responses were obtained from the RNFL

We were able to extract functional assessment of responses carried along RGC axons in addition to those recorded directly from RGC somas. This was possible both in the control eye and the ILM peeled eye, but with double the Z score in ILM peeled eyes relative to the control (figure 4). This could be explained by the increased number of GCaMP-expressing RGCs in the ILM peeled eye and therefore a larger number of GCaMP expressing axons traversing the field of view. We note that the RNFL trace was dominated by an OFF response that may be explained by the stimulus design. A high photopic mean light level can drive stronger OFF responses than ON. An enlarged, pan retinal stimulus could further increase the measured RNFL response by activating additional RGCs. This finding raises the possibility of RGC responses being contaminated by the overlying RNFL. The Z score per pixel is X fold higher in cell somas relative to the RNFL and soma signals therefore dominate in standard recordings but it is important to note that these effects may more visible in deafferented retina. For this same reason, RGC somas make it difficult to extract signals from the RNFL unless recordings are made beyond the edge of the ILM peel. Recording directly from the RNFL may be of interest in pre-clinical models of glaucoma or assessing axon regeneration of transplanted ganglion cells *in vivo* ^27^. Future work may focus on segmenting responses from individual axon bundles to better understand organization of functional responses traveling to the optic disc.

### The ocular inflammation associated with viral vector injections in immune suppressed, ILM peeled animals does not prevent functional recording

Inflammation was observed in the weeks following viral vector injection in the eyes scored in this study (Table 1). The duration and extent of inflammation varied between immune suppressed animals but we did not observe a substantial increase in inflammation relative to a previous report from a non-vitrectomized, non-ILM peeled primate eye intravitreally injected with vector AAV2:CAG:GCaMP6s ^28^. Relative to this animal, M2 scored higher in overall inflammatory response while M3 scored lower with both M2 and M3 recovering faster. The present study used a ganglion cell specific promoter SNCG ^29^ instead of the ubiquitous promoter CAG, a different serotype (7m8 instead of AAV2) and GCaMP generation (GCaMP8 instead of GCaMP6) than the previous study, making a direct comparison challenging, but the general similarities suggest that inflammation does not preclude adoption of this technique in a pre-clinical context.

### Surgical outcomes associated with peeling the ILM

This study demonstrates the feasibility of using an ILM peel to expand the imaging area for functional assessment of macaque RGCs, but also provides new insight into outcomes associated with the surgery itself. Consistent with clinical literature, the most obvious consequence of ILM peel is an altered foveal contour (Figure 6) consistent with previously reported increase in central macular thickness ^30^. CMDS and DONFL were also observed, the latter being associated with small pockets of RGC loss and occurring more frequently in temporal retina where the RNFL is thinner ^31^. This result is consistent with immunohistochemical studies of excised ILM, suggesting that RGCs are removed during the ILM peel process ^32^ but these had not previously been visualized at the cellular scale in vivo. Limited pockets of ganglion cell loss may be tolerable given the increases in the recording area, particularly as their locations are clear.

### Alternatives to ILM peeling

Alternative methods to circumvent the restricted viral transduction imposed by the ILM in primates have been suggested including enzymatic digestion of the ILM with a nonspecific protease ^11,33^, photodisruption ^34,35^ or sub-ILM injection ^36^. Enzymatic digestion has the advantage of not requiring surgery, but has not been attempted in NHPs, where the ILM is thicker than in small animal models. As ILM thickness is eccentricity dependent in primates, it is not straightforward to titrate the correct dose, risking substantial retinal damage. Photodisruption of the ILM has been tested successfully in bovine retinal explants ^34^ using indocyanine green and recently this work has been extended to an *in vivo* rabbit model ^37^, but further testing is necessary before it can be applied to NHP. Lastly an alternative surgical strategy is sub-ILM delivery of the viral vector which may yield similar results while keeping more of the structure intact. As the ILM is composed of Muller cell foot plates, its removal has the potential to cause Muller cell dysfunction ^38^. ILM delivery is therefore desirable, but is surgically very challenging given the thickness of the ILM and there is a risk of delivering the viral vector sub-retinally or sub-hyloid, neither of which would target the GCL.

## Conclusions

This study demonstrates not only that the ILM peel approach allows functional recording from a larger area of the GCL but also that the recordings remain stable over time and are comparable to those in eyes with no ILM peel. As such, this technique may be of value for pre-clinical evaluation of retinal function, both for detecting vision loss and assessing the impact of therapeutic interventions. For cell-based therapies where single cell resolution is needed, a larger area to assess retinal function would greatly increase the chance of success given the potential sparse integration and uncertainty of the locus of cell delivery.

## Acknowledgements

The authors would like to thank Qiang Yang and the Penn Vector Core (RRID: SCR_022432) at the Perelman School of Medicine, University of Pennsylvania. Research reported in this publication was supported by the National Eye Institute of the National Institutes of Health under Audacious Goals Initiative funding award No. U24 EY033275 Accelerating photoreceptor replacement therapy with in-vivo cellular imaging of retinal function, P30 EY001319 (core) and F32 EY032318 Foveal ganglion cell function in the living eye. The content is solely the responsibility of the authors and does not necessarily represent the official views of the National Inst. of Health. This study was supported by an Unrestricted Grant to the University of Rochester Department of Ophthalmology from Research to Prevent Blindness.

## Supplementary Materials

### Modified Hackett-McDonald Scale for Ocular Inflammation in NHPs

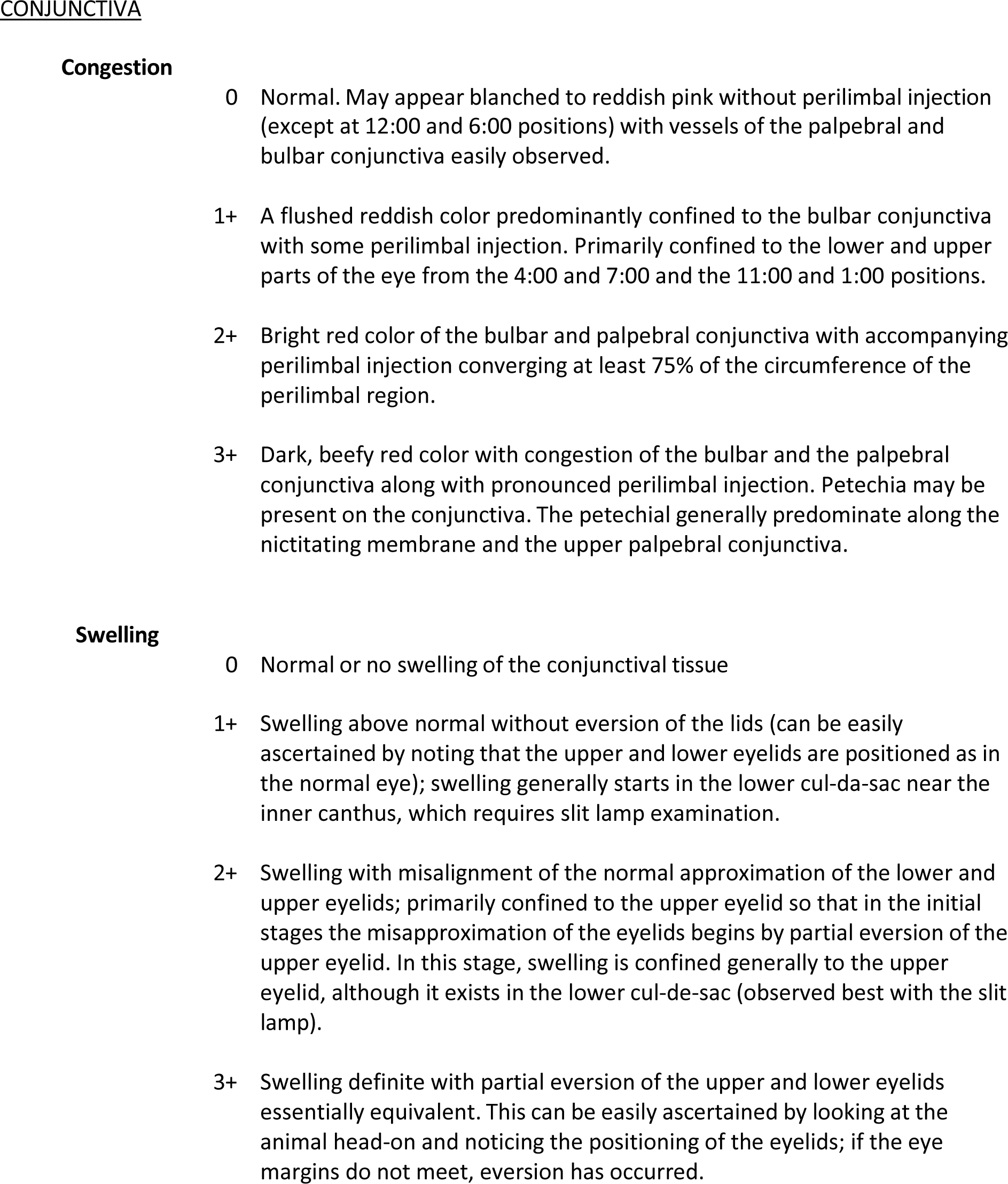
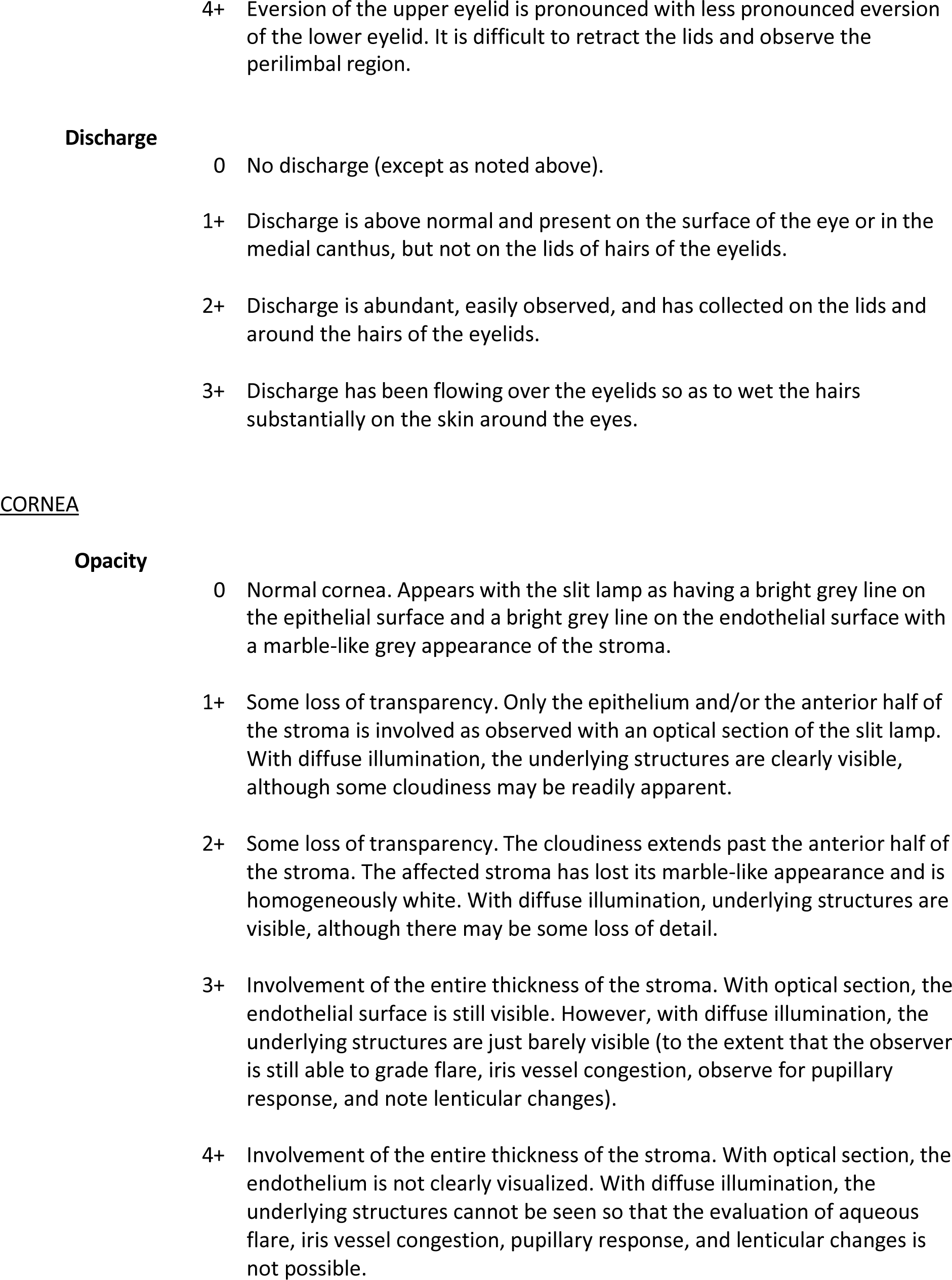
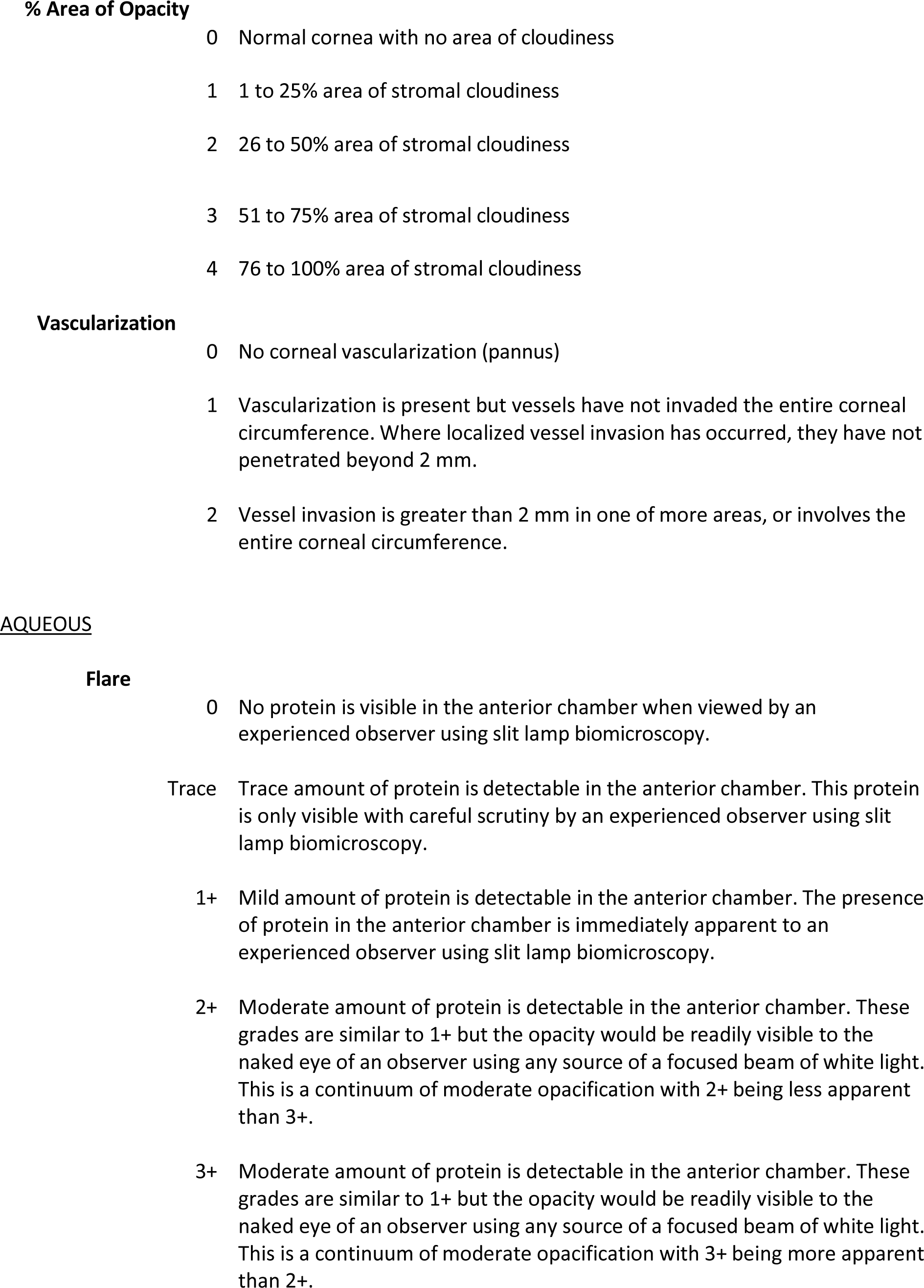
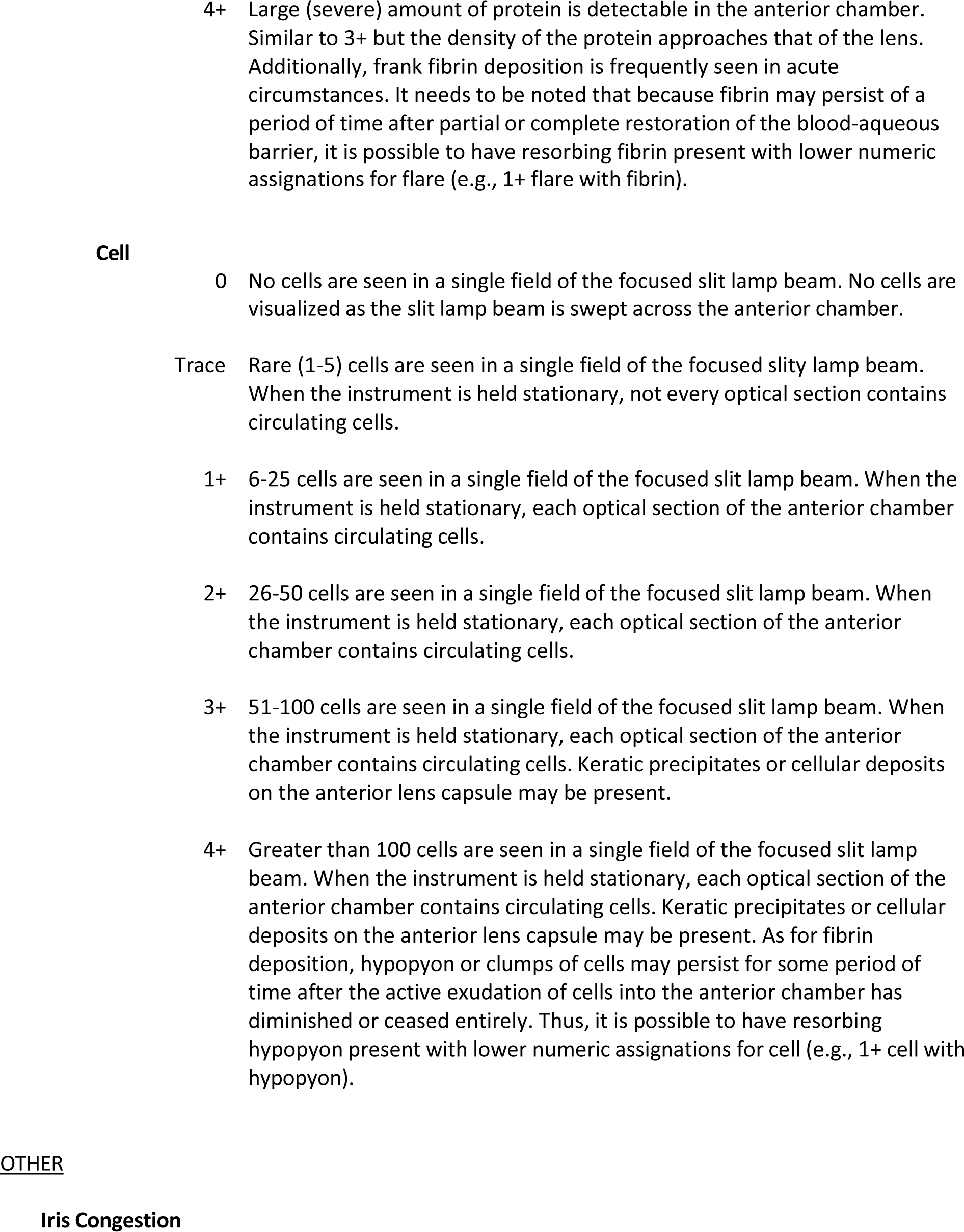
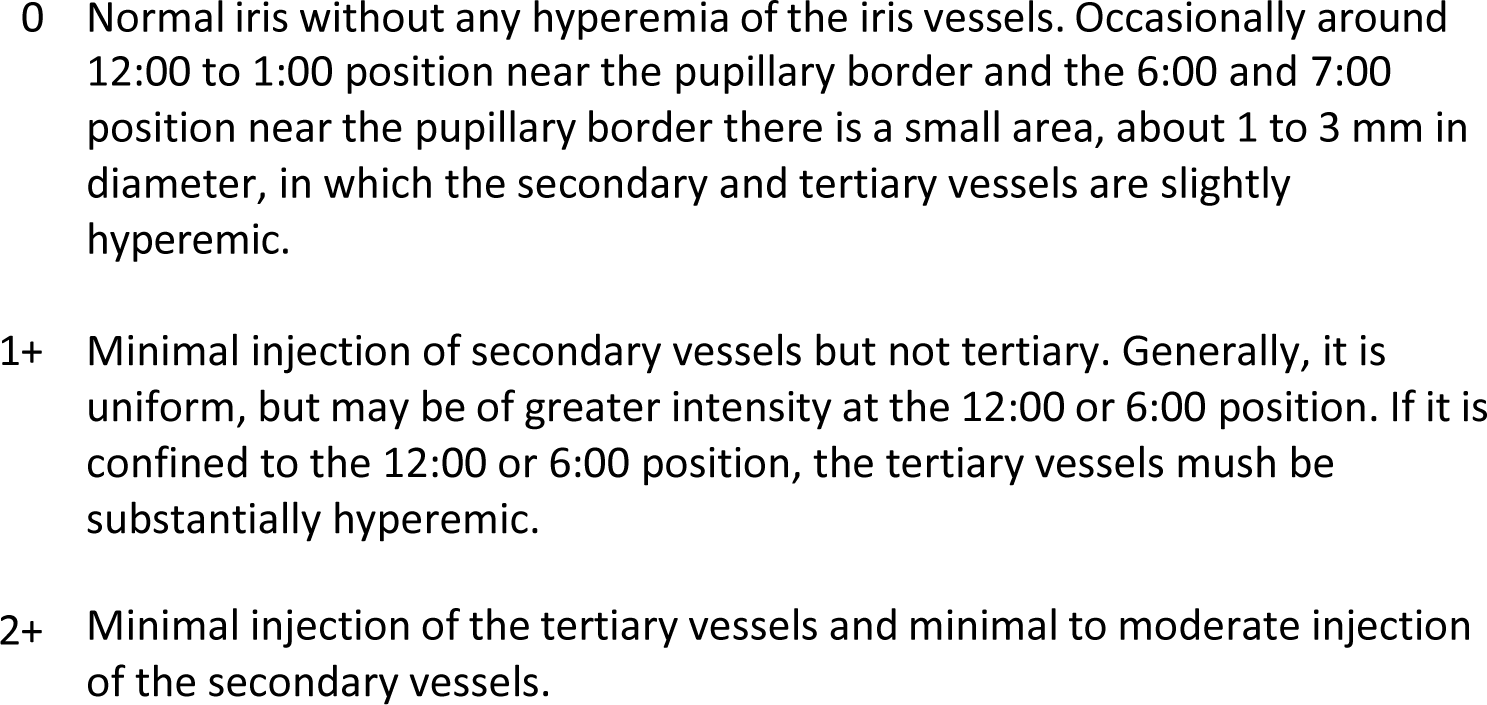
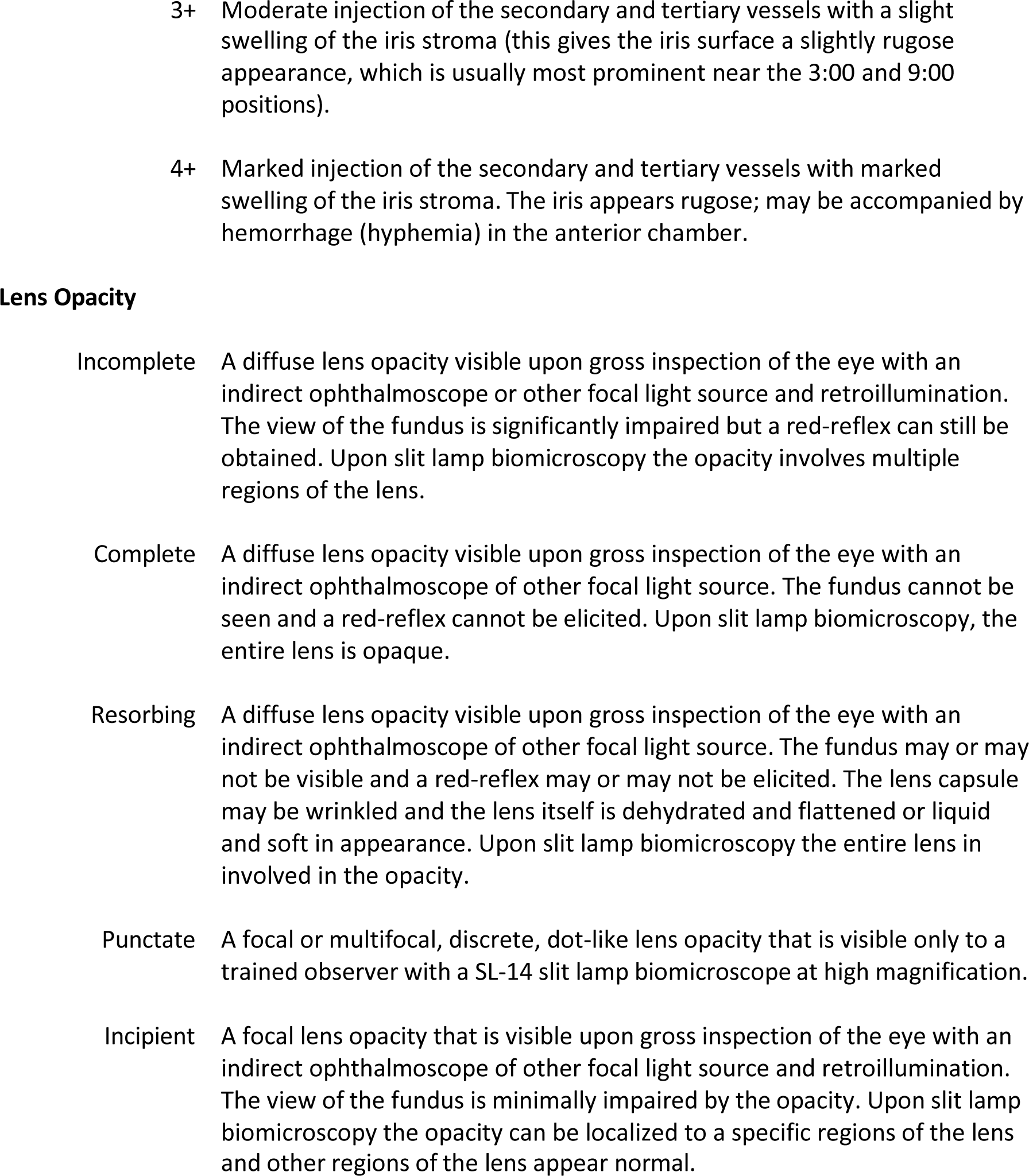

